# Discovering CRISPR-Cas system with self-processing pre-crRNA capability by foundation models

**DOI:** 10.1101/2024.03.11.583506

**Authors:** Wenhui Li, Xianyue Jiang, Wuke Wang, Liya Hou, Runze Cai, Yongqian Li, Qiuxi Gu, Guohui Chuai, Qinchang Chen, Peixiang Ma, Jin Tang, Menghao Guo, Xingxu Huang, Jun Zhang, Qi Liu

**Author notes:** These authors contributed equally (Wenhui Li, Xianyue Jiang).

## Abstract

The discovery and functional annotation of CRISPR-Cas systems laid the groundwork for the development of novel CRISPR-based gene editing tools. Traditional similarity-search-based Cas discovery strategies, which rely heavily on local sequence alignment and reference Cas homologs, may overlook a significant number of remote homologs with limited sequence similarity; and it can not be applied directly for functional recognition. With the rapid development of protein large language models (LLMs), protein foundation models are expected to help model Cas systems with limited Cas homologs without extensive task-specific training data; however, the full potential of these models for Cas discovery and functional annotation has yet to be determined. To this end, we present a novel, effective and unified AI framework, CHOOSER (**<u>C</u>**as **<u>HO</u>**mlog **<u>O</u>**bserving and **<u>SE</u>**lf-processing sc**<u>R</u>**eening), for alignment-free discovery of novel CRISPR-Cas systems with self-processing precursor CRISPR RNA (pre-crRNA) capability utilizing protein foundation models. CHOOSER successfully retrieved 11 novel homologs of Casλ, the majority of which are predicted to be able to self-process pre-crRNA, nearly doubling the current catalog. One of the candidates, EphcCasλ, was subsequently experimentally validated for its ability to self-process pre-crRNA, target DNA cleavage, and trans-cleavage and was shown to be a promising candidate for use as a CRISPR-Cas-based pathogen detection system. Overall, our study provides an unprecedented perspective and methodology for discovering novel CRISPR-Cas systems with specific functions using foundation models, underscoring the potential for transforming newly identified Cas homologs into genetic editing tools.

## 1. Introduction

Clustered regularly interspaced short palindromic repeats (CRISPR)-CRISPR-associated (Cas) systems, which are adaptive immune systems in bacteria and archaea, have been successfully transformed into a variety of powerful genome editing tools over the past decade^1,2^. Ongoing reports of newly discovered CRISPR-Cas systems and subtypes have shown that not only do bacteria and archaea possess these adaptive immune systems, but viruses (particularly phages) also harbor a vast number of undiscovered CRISPR-Cas systems^3–15^.

Among the various DNA-targeting single-effector systems, specific Cas12 effectors, including Cas12a^16^, Cas12i^17^, CasΦ^7^ and Casλ^8^, are known for not only their ability to cleave double-stranded DNA (dsDNA) targets but also their RNase activity, enabling them to process their own precursor CRISPR RNA (pre-crRNA) specifically (**Supplementary Table 1**). These Cas12 effectors can directly utilize compact CRISPR RNA (crRNA) arrays, which are significantly easier to construct than Cas9 single-guide RNA arrays are; this attribute allows the potential use of these effectors for multiplexed genetic modifications, which are crucial for elucidating and modulating gene interactions and networks that underpin complex cellular function. In a previous study, an LbCas12a effector was developed into an effective platform for multiplexed genome engineering that is capable of targeting endogenous genes, with up to 25 individual crRNAs delivered on a single plasmid when a stabilizer tertiary RNA structure is included^18^. Additionally, an engineered variant of Cas12i2, which is guided by a short crRNA without the need for a transactivating crRNA (tracrRNA), has been developed into a versatile and high-performance editing tool for in vivo gene therapy^19^. Given these advancements, functional prioritization of Cas12 candidates with the ability to self-process their pre-crRNA is important because these candidates can enhance the applicability and efficiency of gene editing technologies.

Discovery and functional screening of these CRISPR systems requires the accurate identification of nearby Cas proteins, especially system effectors. Specifically, two main strategies are employed to identify Cas proteins: (1) The first strategy is straightforward that involves identifying Cas homologs based on amino acid sequence similarity. Bioinformatics pipelines, such as NCBI’s Prokaryotic Genome Annotation Pipeline (PGAP)^20^ and CRISPRCasTyper^21^, use this strategy, relying on BLAST alignments^22^ and/or HMM profile searches^23^ against known Cas protein sequences. However, this sequence similarity approach can miss potential distant homologs. A recent study has uncovered 188 rare and previously unrecognized CRISPR-associated gene modules by employing FLSHclust, an algorithm designed for a deep terascale clustering of proteins based on sequence similarity^24^. This study indicates that while Cas proteins demonstrate ‘LEGO-like’ modularity in their sequence arrangements, they maintain conserved key functional domains which can be used for remote homolog searching; however, these alignment-based methods can not be applied directly for functional recognition. (2) The second strategy involves identifying proteins based on structural similarity besides amino acid sequence similarity^4^. This strategy is based on the ‘structure-function’ paradigm prevalent in biology and biochemistry, which states that the functions of proteins are determined by their structures. Therefore, evolving protein structure prediction technologies are expected to play crucial roles in CRISPR-Cas discovery and functional screening. The success of AlphaFold2 demonstrated that its sophisticated transformer-based models can effectively learn relevant representations from multiple sequence alignments (MSAs) of protein homologs, identify evolutionarily conserved patterns and coevolving residues, and subsequently reconstruct the three-dimensional (3D) structure of a protein^25^. In contrast to the two-stage model of AlphaFold2, RoseTTAfold uses rotation- and translation-invariant SE(3) transformer modules in its three-track design, incorporating 1D, 2D, and 3D representations^26^. However, most of these technologies are dependent on a comprehensive reference dataset that includes a sufficient number of known homologs. In the case of Cas protein identification (especially for effectors of some type V and VI subtypes), the number of known homologs is limited, posing a great obstacle for constructing a reference dataset and for the subsequent discovery and functional investigation of Cas proteins using structure prediction tools. Overall, the systematic discovery and functional screening of novel CRISPR-Cas systems with limited homologs and a lack of ground-truth structural information are highly important but challenging.

Large-language models (LLMs) have utilized their powerful representation learning ability in diverse fields, including life sciences. The impressive performance of these foundation models, such as the transformer-based LLM ESM-2, in protein modeling has demonstrated their potential ability to extract and parse essential representations from protein sequences alone, which allows for the reconstruction of protein structures without the need for MSAs^27^. Undeniably, these cutting-edge AI technologies have enhanced our understanding of protein ‘sequence-structure’ connections. Protein foundation models are expected to help model Cas systems with limited Cas homologs for the following advantages: (1) The powerful representation capacity of pre-trained protein LLMs can be leveraged for Cas modeling by fine-tuning without extensive task-specific-training data, where the labeled Cas types are limited in this domain; and the Cas discovery and functional recognition problem can be formulated into a learning-based classification problem as a complement to the traditional alignment-based methods, and (2) the structural characteristics of Cas are expected to be captured by protein LLMs from sequences, thus alleviating the issue of a lack of ground-truth structural information in Cas identification, further facilitating the directly functional recognitions of Cas from sequence alone. However, the full potential of the protein foundation models to address these problems is still undetermined.

To this end, we developed a novel, effective and unified AI framework, named CHOOSER (**<u>C</u>**as **<u>HO</u>**mlog **<u>O</u>**bserving and **<u>SE</u>**lf-processing sc**<u>R</u>**eening), utilizing protein foundation models for alignment-free discovery of novel CRISPR-Cas systems with self-processing pre-crRNA capability. CHOOSER is formulated as a learning-based classification system that can be used to detect potential CRISPR-Cas systems and to directly functionally screen type V Cas12 homologs that possess pre-crRNA self-processing abilities, which is served as a significant complement to the alignment-based methods. In brief, CHOOSER was designed to address the two pivotal questions in CRISPR-Cas system identification and functional screening: (1) An integrative AI strategy was developed to uncover distant Cas homologs by fine-tuning a pretrained large language model, ESM-2. (2) The specific functions of Cas12 enzymes, namely, their ability to self-process pre-crRNA, was directly predicted by leveraging the representations produced by the foundation model. Specifically, CHOOSER successfully identified 3,477 potential CRISPR-Cas systems, enhancing the number of known type II, type V, and type VI systems. Among these, CHOOSER detected 39 Cas12 candidates that had previously been overlooked by the existing alignment-based CRISPR-Cas mining tools, such as CRISPRCasTyper. Of these 39 homologs, 11 novel homologs of Casλ are identified, the majority of which are predicted to be able to self-process pre-crRNA. Subsequent experiments confirmed both the pre-crRNA processing activity and the DNase activity of one of the newly discovered Casλ homologs, EphcCasλ. Overall, our comprehensive study indicated that a proper representation or embedding derived from a protein LLM can be utilized for CRISPR-Cas system identification and functional screening with limited labeled Cas homologs when ground-truth structural information is unavailable. Computational analysis and experimental validation by CHOOSER provide an unprecedented perspective and methodology for discovering novel CRISPR-Cas systems with specific functions using foundation models, underscoring the potential for transforming newly identified Cas homologs into genetic editing tools.

## 2. Results

### 2.1 The framework of CHOOSER

We developed a novel and effective framework, CHOOSER (**<u>C</u>**as **<u>HO</u>**mlog **<u>O</u>**bserving and **<u>SE</u>**lf-processing sc**<u>R</u>**eening), for the discovery and functional screening of CRISPR-Cas homologs with the ability to self-process pre-crRNA via foundation models (**Fig. 1**), which consists of 4 steps:

**Fig. 1.**
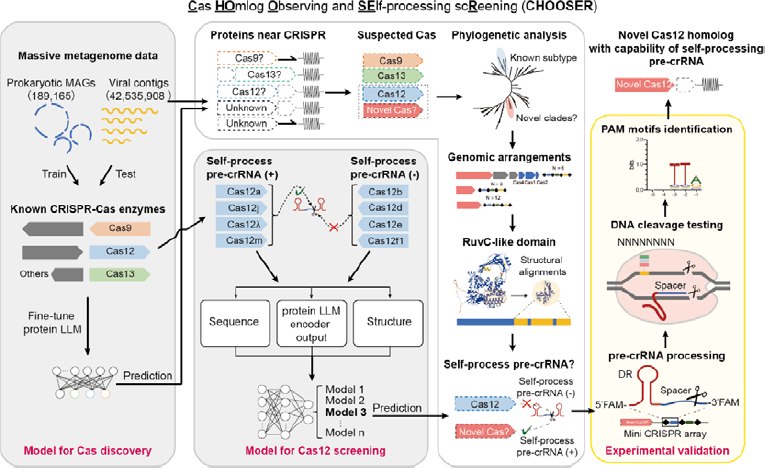
Schematic diagram of the CHOOSER framework for identifying and functional screening of CRISPR-Cas systems with self-processing pre-crRNA capability.

**Step 1. Cas homolog discovery using a protein LLM**. Prokaryotic-origin Cas single-effectors (Cas9, Cas12, and Cas13) were used as training data to fine-tune the protein large-language model ESM-2, and viral-origin homologs were used as testing data to evaluate its performance. The fine-tuned model enables us to identify potential Cas9, Cas12, and Cas13 homologs from other microbial background proteins using protein sequence information alone without extensive task-specific training data.

**Step 2. Pre-crRNA self-processing functional screening of Cas12 homologs using the protein LLM.** We prioritized Cas12 candidates capable of self-processing their pre-crRNA using embedding derived from ESM-2. Our comprehensive tests indicated that the intermediate encoder outputs of ESM-2, as the representations of the models, is all that is necessary for Cas functional screening after examining a variety of representations from sequences to structures.

**Step 3. Phylogenetic analysis and identification of novel candidate Cas12 enzymes.** We first extracted all proteins near the CRISPR arrays that were not annotated as Cas proteins by HMMER. After a preliminary filtering process based on our criteria, these proteins were analyzed using **Step 1** of CHOOSER for Cas discovery, with the aim of identifying potential Cas homologs. These suspected Cas proteins were subsequently used to construct a phylogenetic tree in the context of known Cas proteins to determine their subtypes. For type V Cas12 candidates, proteins from clades of interest underwent additional structure alignments and MSAs to identify their RuvC-like domains and potential nuclease active site residues. Finally, these Cas12 candidates were evaluated using **Step 2** of CHOOSER to predict whether they were capable of self-processing pre-crRNA.

**Step 4. Enzymic activity validation and protospacer adjacent motif (PAM) identification.** To rapidly determine whether a Cas12 candidate possessed pre-crRNA processing and DNA cleavage activities, we expressed and purified the candidate protein and then conducted several in vitro experiments. (1) We designed a mini-CRISPR array by concatenating the direct repeat (DR) sequence with a spacer to test for protein pre-crRNA processing activity. (2) We used a PCR product library constructed with 8 randomized nucleotides upstream of the 5’ end of the target spacer to assess DNA cleavage capability and to identify functional PAMs preferentially targeted for depletion by the protein. Once a candidate’s enzymatic activities were confirmed, the candidate was considered suitable for further engineering and development as a gene editing tool.

### 2.2 Cas homolog discovery using a protein LLM

We first constructed the training and testing datasets for the Cas discovery model. To this end, we collected 189,165 prokaryotic metagenome-assembled genomes (MAGs) and 42,535,908 viral metagenome-assembled contigs. These sequences were processed using the CRISPRCasTyper pipeline to compile a dataset of known Cas proteins. As depicted in **Fig. 2a**, the dataset for prokaryotic-origin proteins consisted of 1,299 Cas9 proteins, 722 Cas12 proteins, 136 Cas13 proteins, and 246,282 negative background prokaryotic proteins collected from the CRISPRclass19 dataset^28^. This dataset was split into training and validation subsets at a ratio of 8:2. Given the rarity of Cas single-effector proteins in the overall pool of proteins of microbial origin, our dataset was highly imbalanced. The ratio of the target Cas9, Cas12, and Cas13 classes to the negative background class was approximately 10:5:1:1,811. For the testing data, we collected a viral-origin protein dataset, as shown in **Fig. 2b**. Collectively, our datasets contained 188,741 prokaryotic-origin proteins as training data, 59,698 prokaryotic-origin proteins as validation data and 2,168 viral-origin proteins as testing data (see Methods). Our reasons for training the model using prokaryotic-origin proteins and testing it on viral-origin proteins were as follows: (1) Based on a prior study revealing that phage-encoded CRISPR□Cas systems possess all six known Cas types (I∼VI)^8^, we considered it appropriate to assemble a testing dataset of viral-origin proteins. (2) Given the distinctive phage-specific traits of viral-origin Cas homologs, such as CasΦ, which is a phage-specific protein with less than 7% amino acid identity to other type V Cas12 homologs^7^, our choice of a viral-origin testing dataset is aptly suited for evaluating the model’s performance and generalizability across different species. (3) The discovery of novel phage-specific type V subtypes suggested that viral metagenomics data may have considerable potential as a valuable reservoir for identifying functional enzymes.

**Fig. 2.**
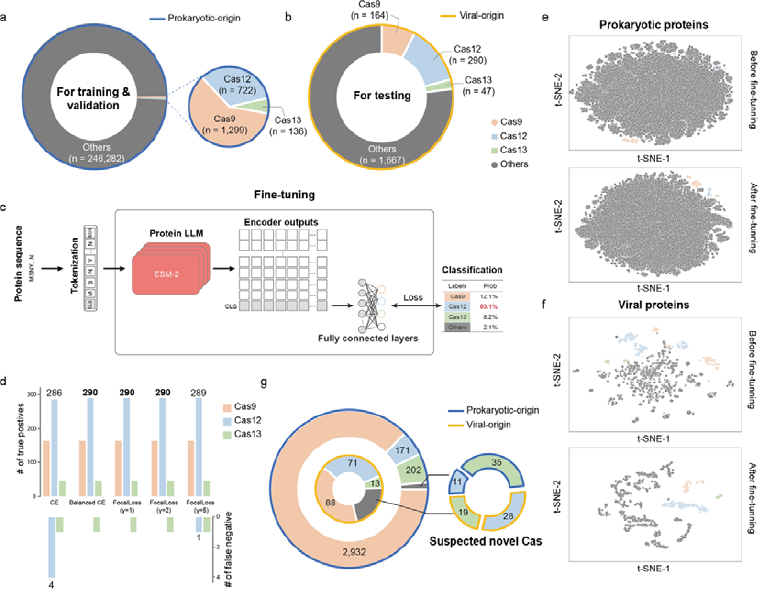
Model trained for Cas single-effector discovery. **a**, Prokaryotic-origin Cas single effectors and other background proteins used as training and validation datasets. **b,** Viral-origin Cas single effectors and other viral proteins used as a testing dataset. **c**, Schematic of the fine-tuned ESM-2 model for discovering Cas single effectors. **d,** Adjustment of hyperparameters for the focal loss to enhance the performance of the classification model. **e-f,** Data distributions for prokaryotic-origin (**e**) and viral-origin (**f**) datasets visualized using the representations extracted by the ESM-2 models before and after fine-tuning. **g**, Suspected Cas homologs identified by our fine-tuned model.

We then built an ESM-2-based Cas homolog classification model and compared it with a baseline sequence model, i.e., the long short-term memory (LSTM) network-based model. In this case, Cas homolog discovery was formulated as a supervised multi-class classification task. This task was designed to differentiate homologs of Cas9, Cas12, and Cas13 from a pool of other proteins of microbial origin. For the baseline LSTM model, each protein’s amino acid sequence was padded to a uniform length of 1,560 and then tokenized to serve as inputs for the models. We observed that the bidirectional LSTM (BiLSTM) model, despite being configured with an embedding size of 1024, a hidden size of 512, and 36 network layers, was unable to meet the requirements of our multi-class classification task for the highly imbalanced datasets. This inadequacy persisted even after the application of a focal loss function during the training process. These results suggested that conventional language model architectures, such as LSTM, may lack the necessary representation capacity to extract critical features from relatively long protein sequences and severely imbalanced data. On the other hand, in the protein LLM ESM-2, taking advantages of the fine-tuning strategy, we utilized the network’s final layer of encoder output corresponding to the CLS token as a representation for each protein. These representations were then connected to fully connected layers to execute the classification task, as depicted in **Fig. 2c** (see Methods).

Considering the extreme imbalance of our training datasets, we experimented with several loss functions, including balanced cross entropy (CE) loss and focal loss, and adjusted the hyperparameters in our multi-class classification task (see Methods). As depicted in **Fig. 2d**, when fine-tuned with protein sequences of prokaryotic origin, the test results demonstrated that the pre-trained protein LLM is effective for “few-shot” identification of the distant Cas proteins such as the Cas12i, using the highly imbalanced training data, which only has 3 training samples for this specific Cas12 subtype. Also it holds the capacity of “zero-shot” identification of viral-specific Cas12 subtypes, i.e, the viral-specific CasΦ, where this subtype does not exist in the training data. Notably, the optimized models successfully detected all viral-origin Cas12 homologs while utilizing a balanced CE loss function. These results clearly demonstrated the effective representation capacity of protein foundation models in Cas homologs discovery. Detailed model performance metrics are provided in **Supplementary Table 2**.

To verify the effectiveness of the ESM-2 model in capturing essential features for distinguishing Cas single-effectors, we visualized the high-dimensional encoder output embeddings from our prokaryotic-origin and viral-origin datasets using Principal Component Analysis (PCA) and t-distributed Stochastic Neighbor Embedding (t-SNE). As depicted in **Fig. 2e and f**, after fine-tuning the embeddings produced by ESM-2, the model was capable of clearly segregating Cas single effectors from the background proteins of both prokaryotic and viral origins. An expanded visualization of the data distribution of our datasets utilizing representations from the fine-tuned ESM-2 models with varying hyperparameters for the loss function is provided in **Supplementary** Fig. 1. These results indicated that the fine-tuned pretrained protein language models effectively extracted the essential features necessary to distinguish Cas single effectors.

In summary, in this study, we utilized a fine-tuned ESM-2 model using balanced CE loss to detect suspected Cas single effectors from all suspected proteins near CRISPR arrays. Using our fine-tuned model, we identified putative 3,020 type II, 242 type V, and 215 type VI systems that were not reported by alignment-based CRISPRCasTyper. In addition, our model discerned 46 and 47 potential novel Cas single effectors from prokaryotic and viral data sources, respectively (**Fig. 2g**). The newly discovered proteins were considered suspected Cas homologs in subsequent analyses.

### 2.3 Functional screening of Cas12 homologs with self-processing pre-crRNA capability using the protein LLM

#### Cas12 training data curation and pseudolabeling

In this study, we aimed to establish an in silico strategy to predict whether Cas12 homologs possess the self-processing pre-crRNA functional trait, and this task is formulated as a binary classfication problem. The premise to achive this goal is to functionally label the distinct known Cas12 homologs as the training data for model building. As more subtypes of type V systems have been successively discovered, the nuclease-associated properties of diverse Cas12 homologs have been revealed. As summarized in **Supplementary Table 1**, certain Cas12 effectors utilize a variety of mechanisms for pre-crRNA processing. To this end, we gathered known Cas12 homologs, with duplicates removed, to construct the training and validation datasets. Cas12a, Cas12c, Cas12i, CasΦ, and Cas12m homologs were labeled pseudopositives for self-processing pre-crRNA, while the remaining Cas12 subtypes were designated as pseudonegatives, as described in the literature and illustrated in **Supplementary Table 1**. Notably, the newly published Casλ (pseudopositive) and Cas12n (pseudonegative) homologs were excluded during the training process. For the testing dataset, we utilized recently published sequences from the CasPEDIA database^29^, with duplicates eliminated with a sequence similarity less than 70% compared to the training dataset. This included several Casλ and Cas12n homologs. These datasets were curated for subsequent experiments designed to probe the functional traits of Cas12 proteins (see Methods).

#### Embedding derived from LLMs is likely all you need for self-processing pre-crRNA functional screening of Cas12 homologs

Guided by the ‘sequence-structure-function’ paradigm, we explored a variety of representations, ranging from sequence to structure, as input for the classifiers to distinguish proteins with unique self-processing pre-crRNA functional properties. Considering the lack of ground-truth structural data for most known Cas proteins and our potential novel candidates, we first evaluated the reliability of state-of-the-art folding structure prediction tools, such as AlphaFold2, RoseTTAfold2 and ESMfold, in predicting the structures of Cas12 homologs (see Methods and Supplementary Information). The ESMfold-predicted structures were fairly consistent, and ESMfold outperformed AlphaFold2 and RoseTTAfold2 in both structural alignment and distance map identity (**Fig. 3b and c**). This could be attributed to the ability of ESMfold to model Cas structures without requiring MSAs from homologs. Therefore, given the aim of our study to identify and screen potential novel CRISPR-Cas systems, we decided to employ ESMfold-predicted structures to derive our subsequent structure embedding.

**Fig. 3.**
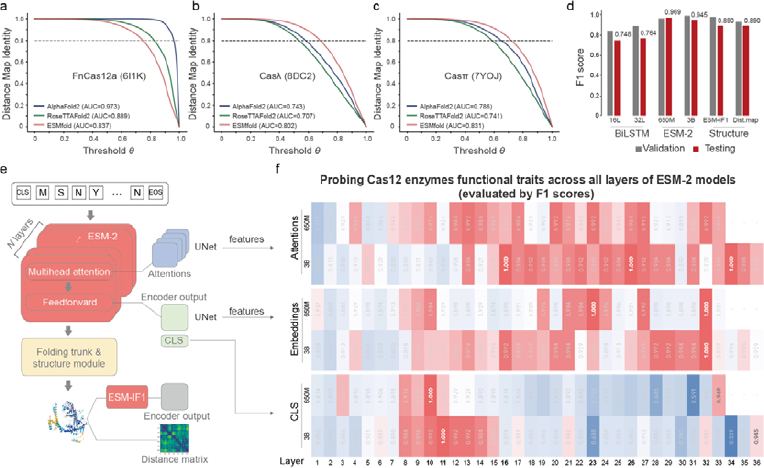
Model trained for predicting Cas12 enzymes capable of self-processing pre-crRNA. **a-c**, Curves of identity between Cα distance maps from cryo-EM structures and their corresponding predicted protein structures, along a range of thresholds : **a,** FnCas12a (PDB ID: 6I1K); **b,** Casλ (PDB ID: 8DC2); and **c,** Casπ (PDB ID: 7YOJ). **d,** Performance of the models in predicting the ability of Cas12 proteins to self-process their pre-crRNA, denoted by F1 scores in the validation (in gray) and testing (in red) datasets. **e,** Assorted representations from protein sequences to structures throughout the ESMfold protein structure prediction process. **f,** F1 scores on the testing dataset, illustrating the performance of the models in predicting the ability of Cas12 protein self-processing pre-crRNA, using varied representations.

We then assessed whether the representation of amino acid sequences and folding structures could discern the unique trait of self-processing of pre-crRNA in Cas12 homologs (see Methods). For the amino acid sequences, two sequence embeddings were applied: (1) we encoded the whole protein sequences into a fixed-length vector as a representation to feed into the bidirectional LSTM (BiLSTM) networks, configured with 16 layers and a hidden size of 128 (16L), as well as 32 layers and a hidden size of 512 (32L); (2) we used the last hidden layer outputs of protein CLS tokens of the transformer-based pretrained protein LLM ESM-2 models, with parameter sizes of 650 million (650M) and 3 billion (3B), for classification. For the folding structures, we adopted two distinct data processing approaches: (1) inverting the folding structures into embedding using the cutting-edge geometric vector perceptron (GVP)-transformer pretrained model ESM-IF1^30^ and (2) converting the three-dimensional folding structures into two-dimensional distance map matrices. We subsequently deployed a convolutional network, U-Net^31^, to extract features from the inverse folding embedding and distance maps, further facilitating classification (see Methods).

The F1 scores from classifications using various input embeddings in the testing dataset are displayed in **Fig. 3d**. Significantly, classifications using ESM-2 embedding significantly outperformed both the BiLSTM models and the structural methods. In detail, the BiLSTM models demonstrated relatively high recall but very low precision. For instance, the BiLSTM 16L model successfully identified 10 out of 12 Casλ homologs as positives but incorrectly predicted 18 out of 40 pseudonegative Cas12n homologs as positives. In contrast, the structural methods displayed better precision but significantly lower recall. For example, ESM-IF1 embedding identified only one Casλ homolog as positive for self-processing pre-crRNA, failing to distinguish the other 11 Casλ homologs. With the ESM-2 embedding methods, we observed both high recall and high precision. Specifically, classification with ESM-2 650M CLS token representations achieved a significantly high recall of 1.0, although with slightly lower precision, while classification using ESM-2 3B CLS token representations achieved a high precision of 1.0, although with slightly lower recall. More details can be found in **Supplementary Table 4**.

Collectively, our comprehensive tests indicated that the embedding derived from LLMs may be all that is necessary for self-processing functional screening of Cas12 homologs when ground-truth structural information is unavailable.

#### Interpretability analysis of the ESM-2 model for self-processing pre-crRNA functional screening

Considering that previous research has demonstrated that transformer-based language models can extract the structural and even functional properties of proteins using attention mechanisms^32^, we sought to delve more deeply into the interpretability of the hidden layers of ESM-2 to determine where the identification of Cas12 homologs capable of self-processing pre-crRNA is encapsulated. As depicted in **Fig. 3e**, ESM-2 adopts a BERT-style encoder-only architecture, and each encoder layer within the ESM-2 model comprises a multihead attention sublayer and a feedforward sublayer. We scrutinized each layer in the ESM-2 models by employing U-Net architectures to extract interpretable features from the encoder output embeddings and the attention weights (see Methods). The CLS token representations of each encoder layer output were also used as input to a classifier tasked with predicting the ability of the Cas12 enzyme to self-process pre-crRNA as previously described. In a manner analogous to the previous experiment, the F1 scores of the probing classifier in our testing dataset served as an indication of the knowledge of this property encoded in the representations. As shown in **Fig. 3f**, both the 650M and 3B models demonstrated that deeper layers of encoder embeddings and attention weights yielded F1 scores reaching 1.0 in the probing classifier. When used as inputs to the classifier, the CLS token representations, which is the first row of the encoder embedding, also attained F1 scores of 1.0 in the 10^th^ and 11^th^ layers of the 650M and 3B models, respectively. These results outperformed the scores obtained using the CLS token representations of the final layers. Taken together, these results indicate that particular properties, in this case, the functional traits of Cas12 homologs, may be present within the intermediate layers of the model rather than within the final layer.

### 2.4 Phylogenetic analysis and identification of novel candidate Cas12 enzymes

Using the Cas homolog discovery model of **Step 1** of CHOOSER, we first identified 39 candidates as potentially novel Cas12 homologs that had not been detected by existing CRISPR-Cas discovery tools, such as CRISPRCasTyper. To verify these candidates, we conducted several analytical steps as follows:

1. **Phylogenetic analysis**. The suspected novel Cas12 candidates were clustered into five distinct clades within the CasPEDIA’s type V Cas12 phylogenetic tree, as depicted in **Fig. 4a** (see Methods).
2. **Genomic loci examination**. We examined and illustrated the genomic arrangements of representatives for each clade in **Fig. 4b**.
3. **Sequence alignments**. We aligned the protein sequences of the candidates against the NCBI ‘nr_clustered’ database (updated 2023-12-28) and against the newly published CRISPR associations identified by the FLSHclust algorithm^24^ using the ‘BLASTP’ algorithm (details in **Supplementary Table 8**).
4. **Structural alignments**. We compared the predicted folding structures of the candidates with those of RuvC proteins to determine the presence of RuvC-like domains and potential Asp-Glu-Asp (D-E-D) active site residues within those domains (see Methods).
5. **Pre-crRNA self-processing capability prediction**. Using the self-processing pre-crRNA screening model of **Step 2** of CHOOSER, we estimated the likelihood that each candidate had the ability to self-process pre-crRNA.

**Fig. 4.**
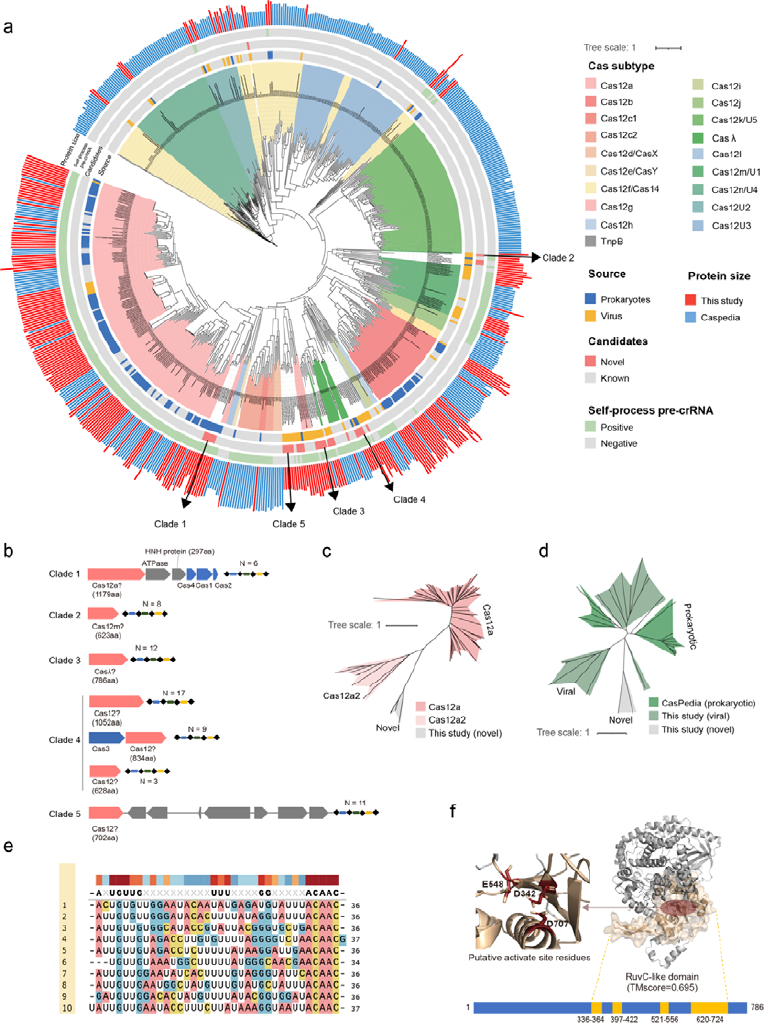
Analysis of 39 novel Cas12 candidates. **a**, Phylogenetic tree of all the suspected Cas12 enzymes against the background of the CasPEDIA type V Cas12 dataset. **b**, Genomic arrangements of representatives of the five clade candidates. **c**, Clade 1 candidates shown phylogenetically against the background of Cas12a homologs. **d**, Clade 2 candidates shown phylogenetically against the background of Cas12m homologs. **e**, MSAs for the direct repeats of Clade 3 suspected CRISPR systems. **f**, Structural alignment showing the RuvC-like domain and canonical D-E-D active site residues within the domain of a putative Casλ homolog in Clade 3.

In detail, clades 1 and 2 were categorized into the Cas12a and Cas12m subtypes, respectively. However, the unique phylogenetic subclusters, as depicted in **Fig. 4c and d**, might account for the fact that the existing Cas detection tools missed them. **Supplementary** Fig. 3 provides an expanded view of these two clades.

The 11 viral-origin proteins categorized within **Clade 3**, with sequence lengths ranging from 735 to 798 amino acids, were phylogenetically proximate to the Casλ clade (**Fig. 4a**). Most of these proteins exhibited a clear CRISPR-Cas system arrangement within their genomic loci, as shown in **Fig. 4b**. Sequence analysis confirmed substantial similarity between all the candidates and the ‘CasLambda’ (PDB ID: 8DC2_A) protein, meeting the significance threshold with e-values of 10^-10^. The CRISPR repeat sequences of these candidates closely mirrored the direct repeat (DR) sequences documented in prior research^8^, as presented in **Fig. 4e**. Structural analysis revealed the RuvC-like domain and the D-E-D catalytic site residues within this domain across the candidates, as shown in **Fig. 4f**. Considering the known pre-crRNA self-processing ability of Casλ enzymes, the prediction that most candidates (9 out of 11) possess this trait is consistent with existing knowledge, suggesting that **Clade 3** significantly enriches the diversity of the Casλ family, nearly doubling its current catalog.

Candidates from clades 4 and 5 did not correspond to any recognized subtypes but established distinct groups. Specifically, **Clade 4** is located phylogenetically between Cas12h and Casλ, while **Clade 5** lies between Casf1 and Cas12d. **Clade 4** candidates varied widely in size, ranging from 628 to 1052 amino acids, and their genomic loci revealed at least three unique arrangement types. **Clade 5** proteins were notably separated from their associated CRISPR arrays within the genome. Sequence comparisons indicated that most proteins in these clades bore significant resemblance to ‘transposase’ proteins. This result raises the possibility that clades 4 and 5 may contain transposases. **Supplementary** Fig. 4 provides an expanded view of these two clades.

### 2.5 Experimental enzymatic activity validation of EphcCas**λ**

For further validation of the newly identified Cas candidates, we focused on EphcCasλ, one of the 11 proteins identified in Clade 3. EphcCasλ, which was discovered in metagenome-assembled contigs from bovine rumen microbial samples, was taxonomically inferred to be akin to ‘Enterobacter phage Phc’. This inference hinted that the enzyme could demonstrate an optimal temperature preference akin to that of mammalian systems. Consequently, EphcCasλ was selected for further experimental validation, with an aim to evaluate its capabilities in pre-crRNA processing and DNA cleavage (see Methods and Supplementary Information).

First, we evaluated the ability of EphcCasλ to self-process its pre-crRNA. Considering that type V CRISPR-Cas systems must generate mature crRNA to facilitate DNA cleavage or interference, we engineered a mini-CRISPR array by fusing a direct repeat sequence with a 20-nucleotide spacer, which allowed for the creation of mini-pre-crRNA. For visualization purposes, we tagged the 5’, 3’, or both ends of the pre-crRNA with 5(6)-carboxyfluorescein (FAM) during the synthesis process, yielding fluorescent mini pre-crRNA, as illustrated in **Fig. 5a** (see Methods). When these fluorescent pre-crRNA substrates were incubated with purified EphcCasλ protein at 37 °C for one hour, we observed efficient cleavage at the 3’ end of the spacer region, as demonstrated by 15% TBE-UREA gel electrophoresis (**Fig. 5b**). This cleavage pattern is consistent with previously described activities of Casλ enzymes.

**Fig. 5.**
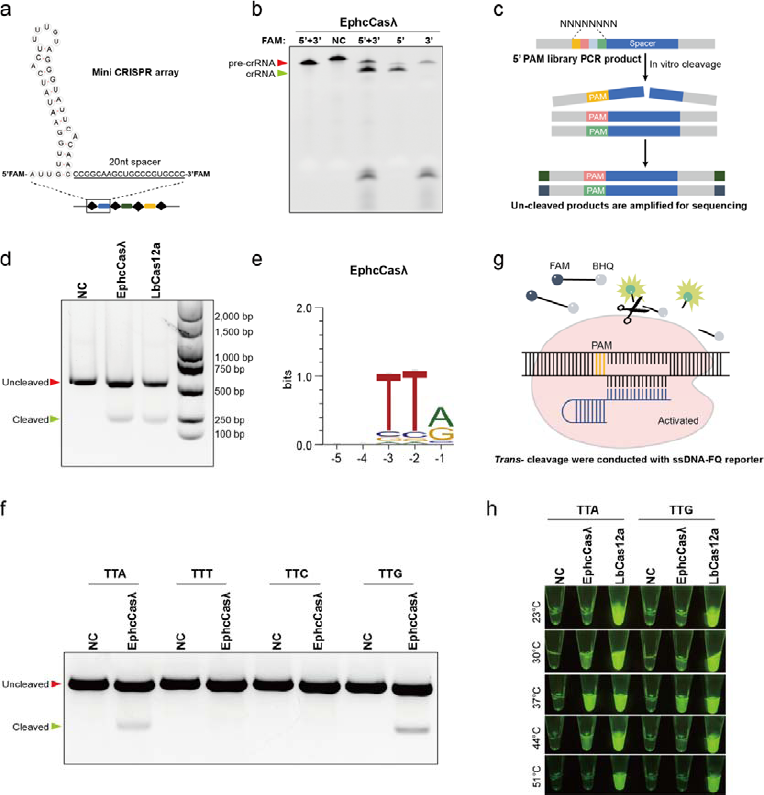
Biochemical characterization of EphcCasλ. **a,** The structure of pre-crRNA with 5’-FAM and/or 3’-FAM were determined via a crRNA processing assay, which included a 36 nt direct repeat sequence and a 20 nt spacer targeting a randomized PAM library. **b,** EphcCasλ specifically processes its own pre-crRNA by cleaving several bases in the spacer region (or 3’ end). A different direct repeat was designed in the NC lane with EphcCasλ RNP complexes, and EphcCasλ could not process this pre-crRNA. **c,** Pipeline used to detect dsDNA cleavage and associated PAM recognition by the EphcCasλ CRISPR system. EphcCasλ RNP complexes cleave a 5’ PAM library PCR product in vitro, and the uncut part was captured via PCR and subjected to Illumina deep sequencing. **d,** EphcCasλ cleaves dsDNA in vitro at 37 °C for 1 h. A 500 bp PCR product was cleaved to two 250 bp products. **e,** Analysis of Illumina deep sequencing data showing that the presumed PAM of EphcCasλ was TTR. Weblogo of the presumed PAM that supported target recognition and cleavage are shown. **f,** EphcCasλ cleaved TTA/TTG dsDNA in vitro at 37 °C for 1 h. PAM was confirmed to be TTA/TTG. **g,** Trans-cleavage assay conducted with the ssDNA-FAM/BHQ reporter. **h,** EphcCasλ exhibited trans-cleavage of DNA at 30 °C, 37 °C, and 44 °C when the PAM was TTA/G.

Next, we assessed the preferred PAM of EphcCasλ. Type V CRISPR-Cas systems target DNA sequences preceded by a 2-to 5-base-pair PAM for self-versus nonself recognition. To determine the PAM preference, we formed a complex of the purified EphcCasλ protein with crRNA and incubated it with PCR products containing a DNA library engineered with 8 randomized nucleotides positioned upstream of the 5’ end of the target sequence. After cleavage, uncut PCR products were isolated and purified through gel extraction, effectively removing functional PAMs from the library (see Methods). The results of PAM depletion analysis confirmed the DNA cleavage activity of EphcCasλ and revealed that it recognizes TTR as its preferred PAM (**Fig. 5c, d, e, and f**).

Furthermore, we also probed the trans-cleavage potential of EphcCasλ on single-stranded DNA (ssDNA). Employing a previously described fluorophore-quencher (FQ) reporter assay^33^, we examined the nonspecific ssDNA cleavage (trans-cleavage) activity of EphcCasλ (see Methods). We observed that the trans-activation of EphcCasλ was most robust at 37 °C, with diminished signals at 30 °C and 44 °C and no activity at 23 °C or 51 °C (**Fig. 5h**). These findings indicated that EphcCasλ exhibits temperature-dependent ssDNA cleavage activity at an optimal temperature of 37 °C, suggesting its potential utility in CRISPR-Cas-based pathogen detection technologies.

## 3. Discussions

The discovery of CRISPR-Cas systems laid the groundwork for the development of CRISPR-Cas-based gene editing technologies. Our research introduces the CHOOSER framework, a novel and effective AI methodology that utilizes protein LLMs, such as ESM-2, to identify and functionally annotate CRISPR-Cas systems. To our knowledge, this study represents the first application of protein foundation models in the discovery and functional screening of CRISPR-Cas systems. We demonstrated the utility of CHOOSER by applying it to Cas12 enzymes and discovered dozens of potential candidates that previous HMMER-based bioinformatics tools, such as CRISPRCasTyper, had missed. These results demonstrated that protein LLM-based approaches can significantly complement traditional sequence alignment methods for novel CRISPR-Cas system identification, and they can be utilized directly for Cas functional annotation leveraging the powerful representation capacity of protein foundation models.

Phylogenetic analysis indicated that the newly identified proteins had marginally longer branch lengths than the known Casλ proteins (**Supplementary** Fig. 5a), and structural prediction alignments revealed relatively low TM scores of approximately 0.5 (**Supplementary** Fig. 5b). These findings suggested rapid evolutionary changes in the source phages, supporting the notion that prokaryotic viral metagenomic data could be a substantial resource for the future discovery of functional enzymes.

Among the Cas12 candidates identified by CHOOSER, we experimentally validated the enzymatic activities of a new Casλ homolog, EphcCasλ. We confirmed its specific pre-crRNA processing and DNA target cleavage capabilities and discovered that its trans-cleavage activity was temperature dependent, with an optimal temperature of 37 °C. These findings suggested that EphcCasλ is a promising candidate for further development in CRISPR-Cas-based pathogen detection systems.

Subsequent potential studies based on our results include the following: (1) Although this study initially explored the interpretability of the self-processing pre-crRNA trait of Cas12 enzymes using protein LLMs, the underlying biological mechanisms involved have not been fully elucidated. For instance, the specifics of how the Cas12m homolog processes its own pre-crRNA are unclear^34^. Moving forward, we aim to delve into the causal links between domain-level and residue-specific functionalities, which could provide a theoretical basis for the functional modification of Cas12 enzymes. (2) We preliminarily validated the self-processing pre-crRNA and DNA cleavage capabilities of EphcCasλ, indicating its promise as a candidate for multiplexed genome editing applications. A thorough investigation of the full characteristics of the EphcCasλ enzyme, such as its cleavage activity in mammalian cells, will be performed in the future. Further engineering modifications to develop this enzyme into an effective CRISPR-Cas-based gene editing tool are expected. (3) This study, through the discovery of 11 variants of the Casλ family, revealed that viral-origin Cas proteins may be undergoing rapid evolution; viral genomes mutate swiftly and are subject to natural selection. This finding provides insight into how enzymes from viral sources could serve as excellent gene editing tools to explore in the future. (4) Our findings aligned with recent research showing that protein LLMs can discern functions of prokaryotic viral proteins^35^. Moving forward, we would investigate the potential of CHOOSER for other functional discrimination.

## 4. Methods

### 4.1 Cas homolog discovery using a protein LLM

#### Known CRISPR-Cas system identification

All the prokaryotic metagenome-assembled genomes and contigs used for CRISPR-Cas system mining were obtained from a multitude of public databases. These included MGnify^36^, GMBC^37^, GEM^38^, 4D-SZ^39^, Glacier Microbiomes^40^, IMG/VR (v3)^41^, MGV^42^, Human Virome Database (HuVirDB)^43^, Gut Phage Database^44^, INPHARED PHAge Reference Database^45^, Virus-Host DB^46^, PLSDB^47^, and Bacteriophage (ftp://ftp.sanger.ac.uk/pub/pathogens/Phage/)^48^.

The CRISPR-Cas mining pipeline CRISPRCasTyper (https://github.com/Russel88/CRISPRCasTyper/releases/tag/v1.8.0)^21^ was utilized to detect known CRISPR-Cas systems since this pipeline annotates Cas proteins using the most recently updated Cas HMM profile database, and it has established a reasonable criterion for identifying known CRISPR-Cas systems with high confidence.

#### Training and testing dataset curation for building the Cas discovery model

In total, we identified 6,639 Cas9 homologs, 2, 342 Cas12 homologs, and 485 Cas13 homologs of prokaryotic origin. After removing redundant proteins with a sequence similarity of no more than 70%, we obtained 1, 299 Cas9, 722 Cas12, and 136 Cas13 nonredundant proteins to serve as the training and validation datasets. The proteins other than the Class 2 Cas single effectors in the prokaryotic published genome-assembled data of *Kira S. Makarova* 2020 (ftp://ftp.ncbi.nih.gov/pub/wolf/_suppl/CRISPRclass19/)^28^, a total of 246, 282 proteins after removal of duplicates, served as the negative background proteins during our Cas discovery model training process. To evaluate our models, we combined nonredundant viral-origin proteins, including 164 Cas9 homologs, 290 Cas12 homologs, 47 Cas13 homologs and 1, 667 negative background proteins, to construct the testing dataset. The published CasΦ (Cas12j) proteins that were not used for training our model are included in the test dataset to test whether the models can discern novel subtypes of Cas single effectors.

#### Obtaining suspected proteins near CRISPR arrays

To discover potential unrevealed CRISPR-Cas systems, we gathered all uncharacterized proteins located in proximity to CRISPR arrays from metagenome-assembled datasets, and we excluded those proteins that were already classified as part of known CRISPR-Cas systems by CRISPRCasTyper. CRISPR arrays were identified using MinCED v0.4.2 (https://github.com/ctSkennerton/minced^49^). Our dataset included proteins from the putative CRISPR-Cas loci reported by CRISPRCasTyper, which included single *cas* genes adjacent to a CRISPR array that were identified by HMMER but failed to meet the filtering criteria^21^. Additionally, we encountered certain uncharacterized large proteins (greater than 600 amino acids) adjacent to CRISPR arrays that lacked recognizable Cas homologs both upstream and downstream, suggesting that they might be components of undiscovered CRISPR-Cas systems missed by current detection methods. We combined all these suspected proteins to discover potential novel Cas systems.

#### Fine-tuning ESM-2 for Cas discovery

With the aim of discovering potential novel Cas single effectors, we utilized the pretrained ESM-2 model (esm2_t33_650M_UR50D, https://github.com/facebookresearch/esm^27^) and fine-tuned it for a multi-class classification task. This task involved classifying Cas9s, Cas12s, Cas13s, and background proteins that are not single effectors. We employed the EsmForSequenceClassification class developed by HuggingFace (https://github.com/huggingface/transformers), which uses the CLS token embedding output from the last encoder layer of the ESM-2 model to perform protein classification. We fine-tuned all parameters of both the ESM-2 model and the fully connected layers within EsmForSequenceClassification.

#### Balanced cross-entropy loss and focal loss

Considering the extreme imbalance among classes in our training dataset, we evaluated several loss functions in addition to the conventional cross entropy (CE) loss. By multiplying by a weighting factor α, we achieved a balanced CE loss. The weighting factor α is calculated based on the sample number *x_j_* in each of the *N* classes, as shown in **Equation (1)**:

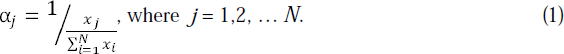

Moreover, we experimented with the focal loss function, incorporating the weighting factor α and applying a range of focusing parameters γ (γ = 1, 2, and 5), as described in a previous research^50^.

#### Embedding visualization

To visualize and compare the embedding representations of the ESM-2 models before and after fine-tuning, we used Principal Component Analysis (PCA) and t-Distributed Stochastic Neighbor Embedding (t-SNE). Initially, PCA was utilized to reduce the dimensionality of the original attribute space of 1,280 dimensions. Then we selected the principal components (PCs) that accounted for 80% of the explained variance for further t-SNE dimensionality reduction and visualization.

### 4.2 Functional screening of Cas12 homologs with self-processing pre-crRNA capability using the protein LLM

#### Training and testing dataset curation for the Cas12 self-processing pre-crRNA functional screening model

For the models for Cas12 screening, we collected all known nonredundant Cas12 proteins and labeled them self-processing pre-crRNA pseudopositives or pseudonegatives according to their subtypes. In detail, 560 pseudopositives (Cas12a, Cas12c, Cas12h, Cas12i, CasΦ and Cas12m homologs) and 345 pseudonegatives (Cas12b, Cas12d, Cas12e, Cas12f, Cas12g, Cas12k, Cas12l and TnpB homologs) were included in the training and validation datasets during the Cas12 screening model training process.

After redundancy removal, the Cas12 homologs published in the CasPEDIA^29^ database included 29 pseudopositives (Cas12a, Cas12c, Cas12h, Cas12i, CasΦ, Cas12m and Casλ homologs) and 98 pseudonegatives (Cas12b, Cas12d, Cas12e, Cas12k and Cas12n homologs), which served as the testing dataset.

#### Evaluating similarities of 3D protein structures

In this study, three state-of-the-art protein structure prediction technologies were employed: AlphaFold2 v.2.1.2, RoseTTAfold2, and ESMfold v.2.0.0. Protein structure alignments were performed using US-align^51^ v.20220626, with template modeling scores (TM-scores) and root mean square deviation (RMSD) used as measurements of structural similarities of pairwise protein structures.

We also calculated the identity of the α-carbon (Cα) distance map matrices as another metric of protein structural similarity. For a protein composed of *n* amino acids, the Cα distance maps of the reference structure and the target structure would be two distinct matrices, denoted as *X* and *Y*. We set a threshold θ _∈_ [0,1] to determine whether each corresponding pair of items, *x_ij_* in *X* and *y_ij_* in *Y*, were in consensus, as defined in Equation (2):

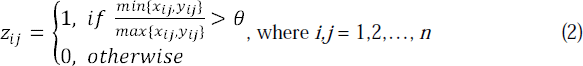

The distance map identity of the protein was subsequently calculated via **Equation (3)**:

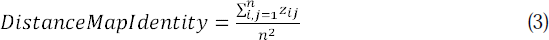

#### Obtaining protein representations extracted by foundation models

For sequence representation extraction, we utilized the ESM-2 pretrained models, specifically esm2_t33_650M_UR50D and esm2_t36_3B_UR50D, to obtain attention weights and output embeddings from each encoder layer, which were subsequently used as inputs for our classifiers.

To generate structural representations, we used the ESMfold-predicted structures in two distinct ways: (1) We used the state-of-the-art geometric vector perceptron (GVP)-transformer pretrained model ESM-IF1^30^ (esm_if1_gvp4_t16_142M_UR50) to transform protein structures into embeddings. (2) We transformed protein structures into two-dimensional distance map matrices using BioPython 1.81 (https://biopython.org/^52^). These structural representations were also used as inputs for our classifiers.

#### Building binary classifiers for functional screening of Cas with self-processing pre-crRNA capability using various representations

For each of the amino acid sequence inputs, we encoded the whole protein sequences into a fixed-length vector as a representation to feed into the bidirectional LSTM (BiLSTM) networks. BiLSTM networks were configured with a hidden size of 128 across 16 hidden layers (denoted as ‘16L’) and a hidden size of 512 across 32 hidden layers (denoted as ‘32L’) to facilitate binary classification tasks. The implementations were executed using PyTorch v1.13.1.

For the amino acid sequence, we also used the ESM-2’s final layer of encoder output corresponding to the CLS token as the LLM-derived representations, followed by the use of fully connected layers for the classification tasks.

Regarding the protein 3D structure inputs, we converted structures into inverse folding embeddings and distance matrices, respectively, as structural representations. Then we applied U-Net networks for feature extraction from these representations and performed binary classifications.

#### Interpretability analysis of the ESM-2 model for self-processing pre-crRNA functional screening

Within each layer of the ESM-2 encoder, we examined three distinct types of representations: (1) the encoder output for the CLS token, (2) the encoder output embeddings, and (3) the multihead attention weights. For the CLS token representation, we utilized fully connected layers for classification as previously described. Regarding the encoder output embeddings and the multihead attention weight matrices, we employed U-Net architectures to extract features, which were then processed through fully connected layers for classification purposes.

### 4.3 Phylogenetic analysis and identification of novel candidate Cas12 enzymes

#### Phylogenetic tree reconstruction

For the phylogenetic analysis of Cas9, Cas12 and Cas13, we used the single effectors newly identified in this study along with homologs provided in the CasPEDIA database. Multi-sequence alignments were generated using MAFFT^53^ v.7.508 with 1,000 iterations and filtered to remove columns composed of gaps in 95% of the sequences. The phylogenetic tree was inferred using IQTREE^54^ v.1.6.12 via automatic model selection and 1,000 bootstraps and visualized using iTOL^55^ v5 (https://itol.embl.de/itol.cgi), as described in previous research^8^.

#### Determination of RuvC-like domains

For the suspected Cas12 candidates, we identified their RuvC-like domains through structural alignment with known RuvC proteins. Specifically, we retrieved the sequences of 522 RuvC proteins from the UniProt^56^ database as of December 12, 2022. We then employed US-align to conduct structural alignments between each Cas12 candidate and the RuvC proteins. The noncontinuous segments of the Cas12 candidates that aligned with any of the RuvC proteins were subsequently identified as RuvC-like domains.

#### Determination of putative active site residues of RuvC-like domains

The suspected Cas12 candidate sequences were aligned with reference Cas12 proteins to identify the conserved active site residues D-E-D within the RuvC-like domains. Multi-sequence alignments using MAFFT were visualized by SnapGene v.6.0.2. Moreover, the spatial conformation of the RuvC-like domains and their conserved active site residues were visualized using PyMOL v.2.5.2.

### 4.4 Experimental enzymatic activity validation of EphcCas**λ**

#### Expression and purification of proteins

EphcCasλ with 6xHis tag overexpression plasmids were transformed into chemically competent *E.coli* BL21 (Vazyme) and incubated overnight at 37°C on LB-Kan agar plate (50 μg/mL Kanamycin, Sangon). Single colony was picked to inoculate 3 ml (LB, 50 μg /mL Kanamycin) starter cultures which were incubated at 37°C for 8 h. Then 500ml 500ml LB-Kan medium(50 μg /mL Kanamycin) were inoculated with 1 ml starter culture and grown at 37°C to an OD600 of 0.8, cooled down on ice and gene expression was subsequently induced with 1 mM IPTG followed by incubation overnight at 16°C. Cells were harvested by centrifugation and resuspended in buffer A (20 mM Tris-HCl, pH 8.0, 500 mM NaCl, 10% Glycerol, 20 mM imidazole), subsequently lysed by sonication, followed by lysate clarification by centrifugation. The soluble fraction was loaded on a 10 ml cobalt column (Thermo Fisher). Bound proteins were washed with wash buffer and subsequently eluted by gradient imidazole (50 mM to 1 M). The eluted proteins were concentrated to 1 mL before loading on a Superdex 200 increase 10/300 GL column (Cytiva) Peak fractions were concentrated to 500 μL and concentrations were determined using a NanoDrop 2000 Spectrophotometer (Thermo Fisher).

#### In vitro pre-crRNA processing assay

6-FAM (Fluorescein)-conjugated pre-crRNA substrates were synthesized by GenScript (substrate sequences are provided in **Supplementary Table 10**). Processing reactions were initiated in a 1:2 molar ratio by combining EphcCasλ protein and RNA substrate in NEB r2.1 buffer and subsequently incubated at 37 °C for 30 min, then cooled down on ice before separation on a 20% TBE-Urea-PAGE. Gels were visualized and imaged by Chemidoc MP imaging system.

#### In vitro PAM depletion assay

A PCR product containing PAM library was amplified from RTW554 plasmid (Addgene, 160132). Active EphcCasλ RNP complex was assembled by mixing protein and crRNAs (GenScript) in a 1:2 molar ratio in NEB r2.1 buffer and incubation at RT for 30 min. In vitro PAM assays were performed in a 20-μL reaction mixture containing 500 ng substrate and RNP complex in a final 1x NEB r2.1 buffer. Assays were allowed to proceed at 37 °C for 2 h. Reactions were then treated with RNase A (NEB) and proteinase K (NEB). Loading dye was added (Gel Loading Dye Purple 6X, NEB) and samples were separated by electrophoresis on a 1% agarose gel. Un-cleaved products were isolated and purified through gel extraction and subsequently amplified by using Phanta Max Super-Fidelity DNA Polymerae (Vazyme) for 25 cycle at the first round PCR (PCR1). And then the products of PCR1 were purified by gel extraction for the second round of PCR (PCR2). For PCR2, DNA was amplified with VAHTS^TM^ DNA adapters set1 for Illumina (Vazyme) using KAPA HiFi HotStart ReadyMix (Roche) for 6 cycles (crRNA and primers sequences are given in **Supplementary Table 10**). After that, the products were purified using DNA Clean Beads (Vazyme) and sequenced on the Illumina Novaseq 6000 platform.

#### PAM depletion sequencing data analysis

Amplicon sequencing of the targeted PCR product was used to identify PAM motifs that were preferentially depleted. Paired-end sequencing reads were merged using FLASH^57^ v.1.2.11 with default parameters. Subsequently, the merged reads were processed as described in a previous study^58^, and the frequency of each possible 8-nucleotide sequence was calculated. Enriched PAMs were determined by calculating the log2 ratio of the abundance of PAMs relative to that of the control plasmids. These enriched PAMs were subsequently used to generate sequence logos.

#### In vitro DNA cleavage assay

The dsDNA substrates were produced by PCR amplification of pUC57 plasmids. Target cleavage assays were performed in a 20-μL reaction mixture containing 300 ng substrate and RNP complex in a final 1x NEB r2.1 buffer (target and crRNA sequences are given in **Supplementary Table 10**). Assays were allowed to proceed at 37 °C for 1 h and samples were analyzed by electrophoresis on a 2% agarose gel.

#### ssDNA trans cleavage and florescence detection

For the trans cleavage assay, 1:2 molar rations of EphcCasλ and crRNA were premixed with the FQ-Labeled reporter (GenScript) in NEB r2.1 buffer and then distributed in 300-μL PCR tubes for the subsequent experiment. Sample containing the activator dsDNA were added to the reporter system and reactions proceed at different temperatures for 1 h, the florescent signal was detected at an excitation wavelength of 480 nm.

## Data availability

Sequencing data are available on the NCBI Sequence Read Archive under BioProject ID PRJNA1074107 (https://www.ncbi.nlm.nih.gov/sra/PRJNA1074107).

## Code availability

The custom code of CHOOSE, together with the trained models for mining CRISPR-Cas systems and screening Cas12 candidates capable of self-processing pre-crRNA, are available at GitHub repositories.

## Acknowledgements

This work was supported by the National Key Research and Development Program of China (Grant No. 2021YFF1201200, No. 2021YFF1200900, No. 2022YFC2702705), National Natural Science Foundation of China (Grant No. 32341008), Shanghai Shuguang Scholars Project, Shanghai Excellent Academic Leader Project, Shanghai Science and Technology Innovation Action Plan-Key Specialization in Computational Biology and Fundamental Research Funds for the Central Universities, Shanghai Municipal Science and Technology Major Project (Grant No. 2021SHZDZX0100), and Youth Foundation Project of Zhejiang Lab (Grant No. K2023PE0AA02).

## Author contributions

QL, XH and WL conceived the project. WL built the CHOOSER framework. WL, WW, RC, GC, QC, PM, JT and MG analyzed the data. WL, LH and YL built and trained the AI models. JZ, XJ, and QG designed experiments, purified proteins and performed biochemical experiments. QL, WH, XH and JZ wrote this manuscript. This manuscript was reviewed and approved by all co-authors.

