## Supplementary Information for "Discovering CRISPR-Cas system with self-processing pre-crRNA capability by foundation models"

### Table of Contents

**List of Abbreviations**

**Performance evaluation of various loss functions applied in fine-tuning protein LLMs for discovering Cas homologs**

**Evaluation of protein folding tools: AlphaFold2, RoseTTAfold2 and ESMfold**

**Supplementary Fig. 1 |** Data distributions for the training, validation, and testing datasets visualized using the representations extracted by the ESM-2 models before and after fine-tuning.

**Supplementary Fig. 2 |** Curves of identity between Cα distance maps from cryo-EM structures and their corresponding predicted protein structures, along a range of thresholds $\theta$.

**Supplementary Fig. 3 |** Genomic arrangements, predicted protein folding structures, RuvC-like domains, putative active sites, and predicted secondary structures of CRISPR repeats for Clade-1 and Clade-2 candidates.

**Supplementary Fig. 4 |** Genomic arrangements, predicted protein folding structures, RuvC-like domains, putative active sites, and predicted secondary structures of CRISPR repeats for Clade-4 and Clade-5 candidates.

**Supplementary Fig. 5 |** Phylogenetic analysis and structural alignments of the Casλ catalog.

**Supplementary Fig. 6 |** Raw data for Fig. 5b. EphcCasλ specifically processes its own crRNA by cleaving several bases in the spacer region (or 3’ end). Similar experiments were performed with 4 μM RNA (left lane1 to left lane5) and 2 μM RNA (left lane6 to left lane10).

**Supplementary Fig. 7 |** Raw data for Fig. 5d.

**Supplementary Fig. 8 |** Raw data for Fig. 5f.

**Supplementary Fig. 9 |** Protein purification results after SEC.

**Supplementary Table. 1 |** Summary of nuclease-associated traits of Cas12 homologs.

**Supplementary Table. 10 |** RNA, plasmids, and primers.

**List of Abbreviations**

**LLM** large langue model

**CRISPR** Clustered regularly interspaced short palindromic repeats

**Cas** CRISPR-associated

**crRNA** CRISPR RNA

**pre-crRNA** precursor CRISPR RNA

**tracrRNA** transactivating crRNA

**dsDNA** double-stranded DNA

**ssDNA** single-stranded DNA

**PGAP** Prokaryotic Genome Annotation Pipeline

**MSA** multiple sequence alignment

**2D** two-dimensional

**3D** three-dimensional

**PAM** protospacer adjacent motif

**DR** direct repeat

**MAG** metagenome-assembled genome

**LSTM** long short-term memory

**BiLSTM** bidirectional LSTM

**CE** cross entropy

**PCA** principal component analysis

**t-SNE** t-distributed stochastic neighbor embedding

**GVP** geometric vector perceptron

**FAM** 5(6)-carboxyfluorescein

**BHQ** Black Hole Quencher

**FQ** fluorophore-quencher

**SEC** size exclusion chromatography

**Performance evaluation of various loss functions applied in fine-tuning protein LLMs for discovering Cas homologs**

The model trained using CE loss function was used as the baseline. As shown in **Fig. 2d**, all the LLM-based models with various loss functions were successful at detecting Cas9 homologs. In the case of Cas12 homologs, we observed that fine-tuned ESM-2 classification models applying various loss functions all detected previously unseen viral-specific CasΦ homologs. In more detail, with the exception of the model using CE loss, which missed one Cas12b homolog, two Cas12f1 homologs, and one CasΦ homolog, the models using balanced CE loss and focal loss (with focusing parameter $\gamma$ =1, 2) exhibited identical performance, identifying all Cas12 homologs in the testing dataset. Notably, when the focusing parameter $\gamma$ was increased to 5, one Cas12f homolog was missed. For the least common class of Cas13 homologs, the failure of all the models to detect a specific Cas13a homolog should not be attributed to the ineffectiveness of the model. The undetected Cas13 protein was found to be incomplete and contained only 607 amino acids. Collectively, these results demonstrated that the protein LLM, fine-tuned with prokaryotic-origin protein sequences, was capable of discovering distant Cas homologs, including those of viral origin.

**Evaluation of protein folding tools: AlphaFold2, RoseTTAfold2 and ESMfold**

We collected 14 published cryogenic-electron microscopy (cryo-EM) structures of Cas12 homologs from RCSB Protein Data Bank (RSCB PDB, https://www.rcsb.org/)[1], and aligned them with their corresponding predicted structures, using the cutting-edge 3D structural alignment algorithm US-align[2]. When comparing the structures predicted by AlphaFold2 with those determined by cryo-EM, we found impressively high TM-scores for well-represented subtypes. This was particularly true for AsCas12a (PDB ID: 5B43), which achieved a TM-score of up to 0.997. Such common subtypes likely benefit from a wealth of homologous sequences available in public databases, which bolster the MSA process central to AlphaFold2's predictive algorithm. Conversely, for recently identified proteins like Casλ (PDB ID: 8DC2) and Casπ (PDB ID: 7YOJ), which have fewer homologous sequences available for effective MSAs, we observed markedly lower TM-scores, with the lowest being 0.263. A similar pattern was observed when assessing the predictive accuracy of RoseTTAfold2-predicted structures in comparison to their cryo-EM counterparts. On the other hand, when aligning ESMfold-predicted structures with the cryo-EM structures, we observed TM-scores with a relatively smaller range of variation. We obtained higher TM-scores, reaching up to 0.827, for smaller-sized Cas12 proteins such as Cas12f1 (PDB ID: 7C7L), Cas12k (PDB ID: 7N3O), and Cas12m2 (PDB ID: 8HHL). In other situations, TM-scores generally hovered around 0.5. The lowest TMscore, dipping to 0.309, was found in Cas12c2 (PDB ID: 7V93), a protein with a size of 1,218 amino acids. Detail results of the structural alignments were provided in **Supplementary Table. 3**.

Besides, we also designed an algorithm to evaluate the identity of α-carbon (Cα) distance maps between cryo-EM structures and their corresponding predicted protein structures. Curves illustrating the distance map identity between cryo-EM structures and their corresponding predicted protein structures are presented in **Fig. 3a-c** and **Supplementary Fig. 2**, with all the corresponding area under the curve (AUC) values provided in **Supplementary Table. 3**.

Notably, ESMfold-predicted structures showed fairly consistent in AUCs, hovering around 0.8. In the context of newly identified and less common subtypes, such as Casλ (PDB ID: 8DC2) and Casπ (PDB ID: 7YOJ), ESMfold outperformed AlphaFold2 and RoseTTAfold2 in both structure alignments and distance map identity AUCs (**Fig. 3b and c**). This could be attributed to the ESMfold's algorithm developed based on the protein LLM ESM-2 model, which does not rely on MSAs of homologs. Taken together, given the aim of our study to identify and screen potential novel CRISPR-Cas systems, we decided to employ ESMfold-predicted structures for our subsequent analyses.

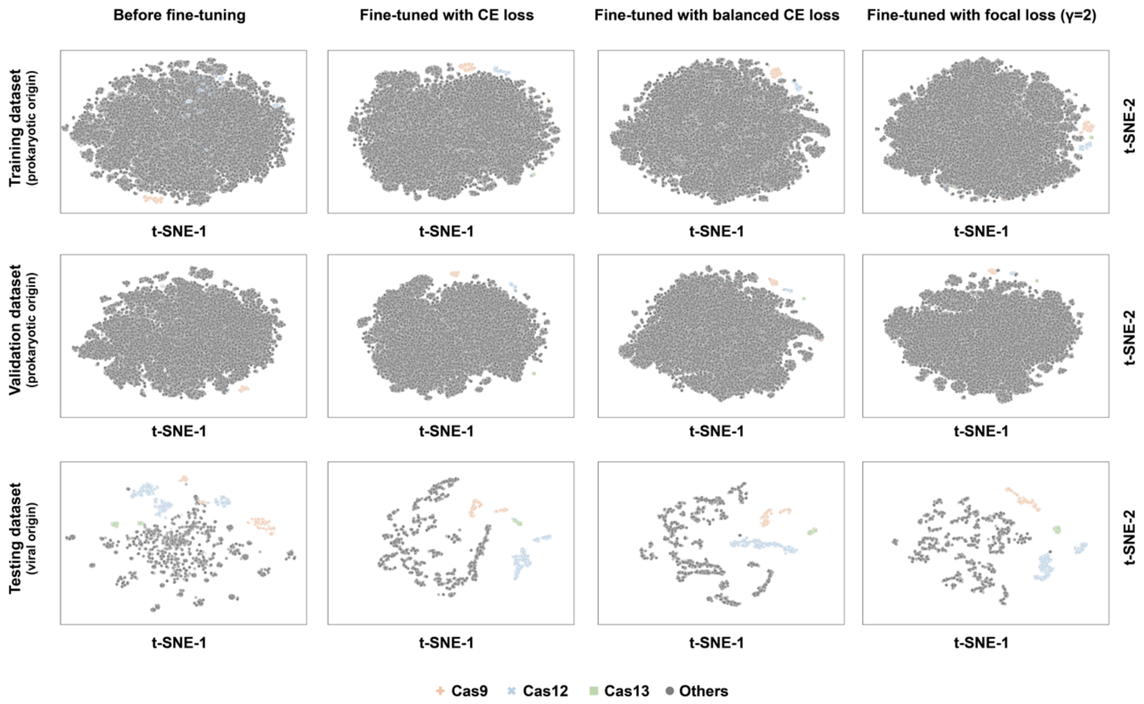


**Supplementary Fig. 1 |** Data distributions for the training, validation, and testing datasets visualized using the representations extracted by the ESM-2 models before and after fine-tuning.

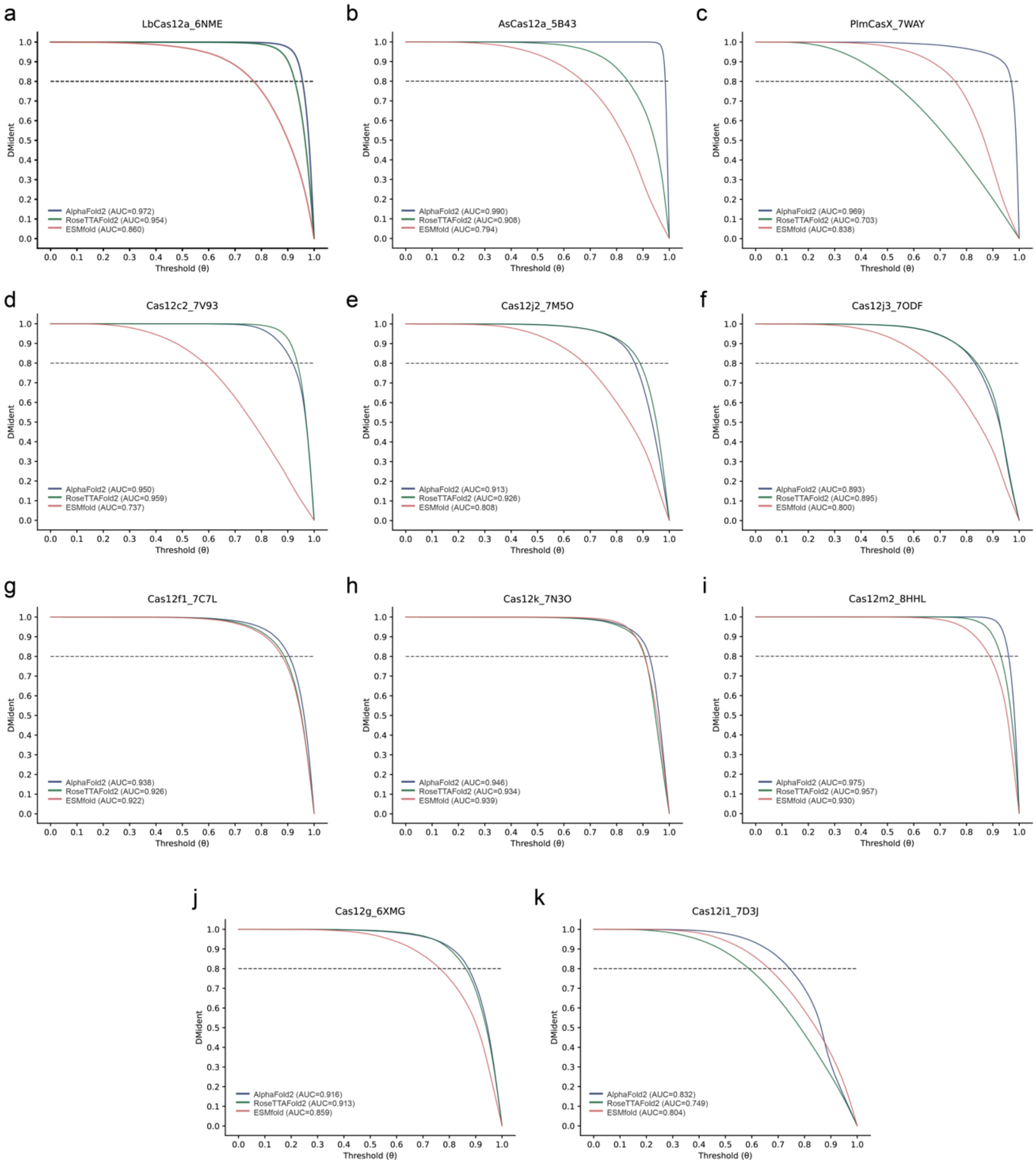


**Supplementary Fig. 2 |** Curves of identity between Cα distance maps from cryo-EM structures and their corresponding predicted protein structures, along a range of thresholds $\theta$: **a,** LbCas12a (6NME), **b,** AsCas12a (5B43), **c,** PlmCasX (7WAY), **d,** Cas12c2 (7V93), **e,** Cas12j2 (7M5O), **f,** Cas12j3 (7ODF), **g,** Cas12f1 (7C7L), **h,** Cas12k (7N3O), **i,** Cas12m2 (8HHL), **j,** Cas12g (6XMG), **k,** Cas12i1 (7D3J).

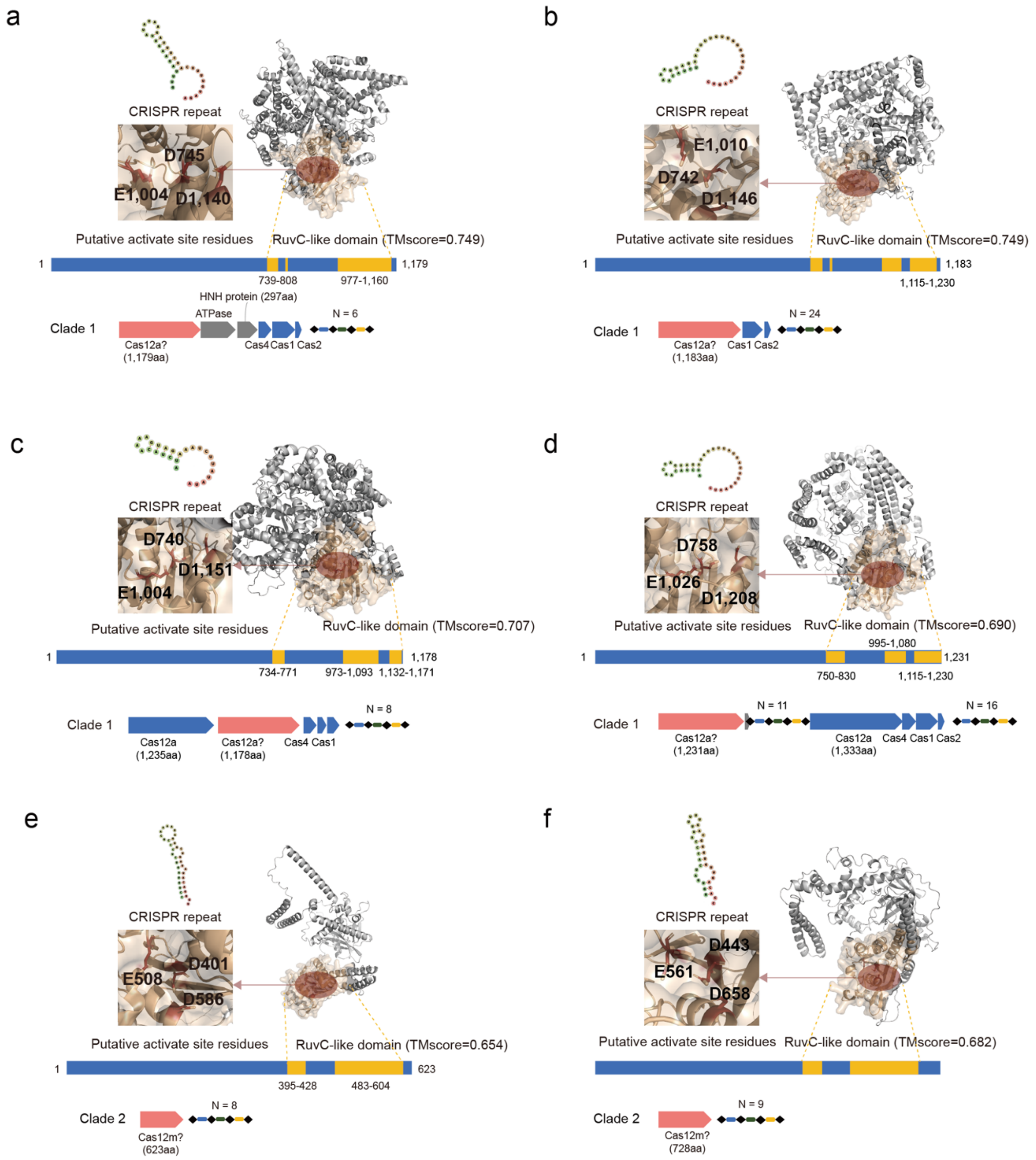


**Supplementary Fig. 3 |** Genomic arrangements, predicted protein folding structures, RuvC-like domains, putative active sites, and predicted secondary structures of CRISPR repeats for Clade 1 and Clade 2 candidates**. a-d,** for Clade 1, putative Cas12a homologs, **e-f**, for Clade 2, putative Cas12m homologs.

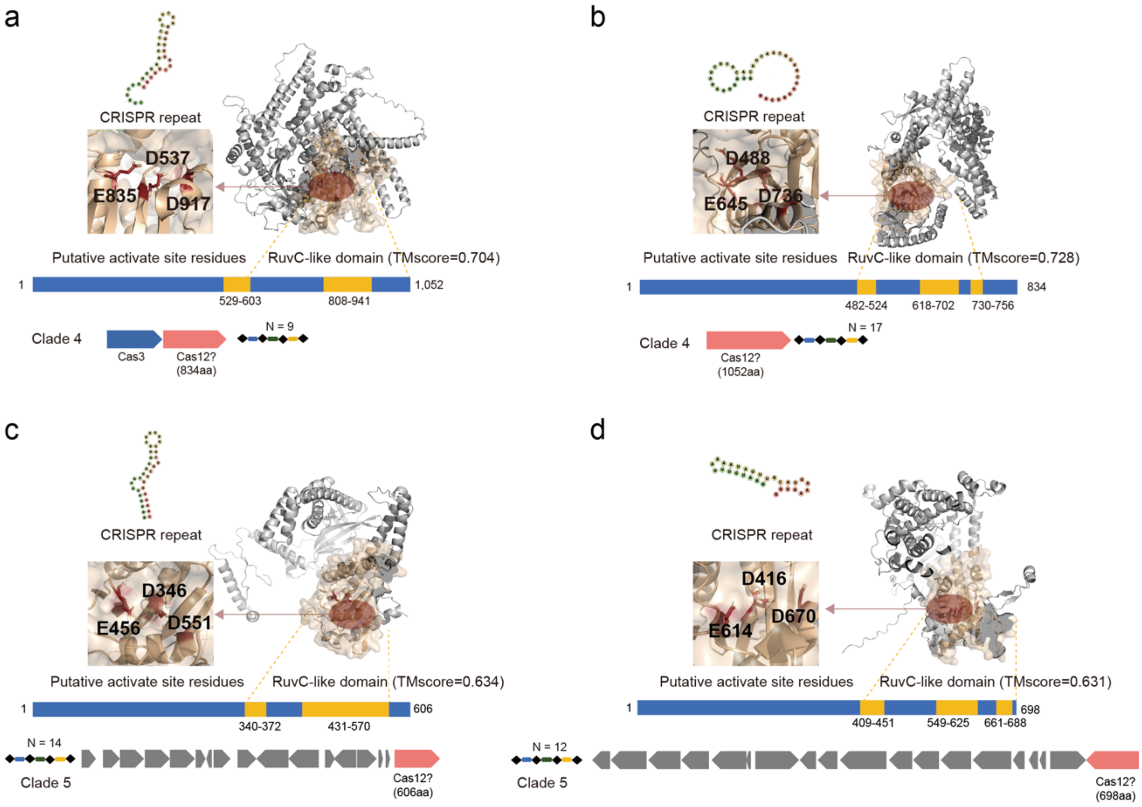


**Supplementary Fig. 4 |** Genomic arrangements, predicted protein folding structures, RuvC-like domains, putative active sites, and predicted secondary structures of CRISPR repeats for Clade 4 and Clade 5 candidates. **a-b,** for Clade 4, putative Cas12 homologs, **c-d**, for Clade 5, putative transposase.

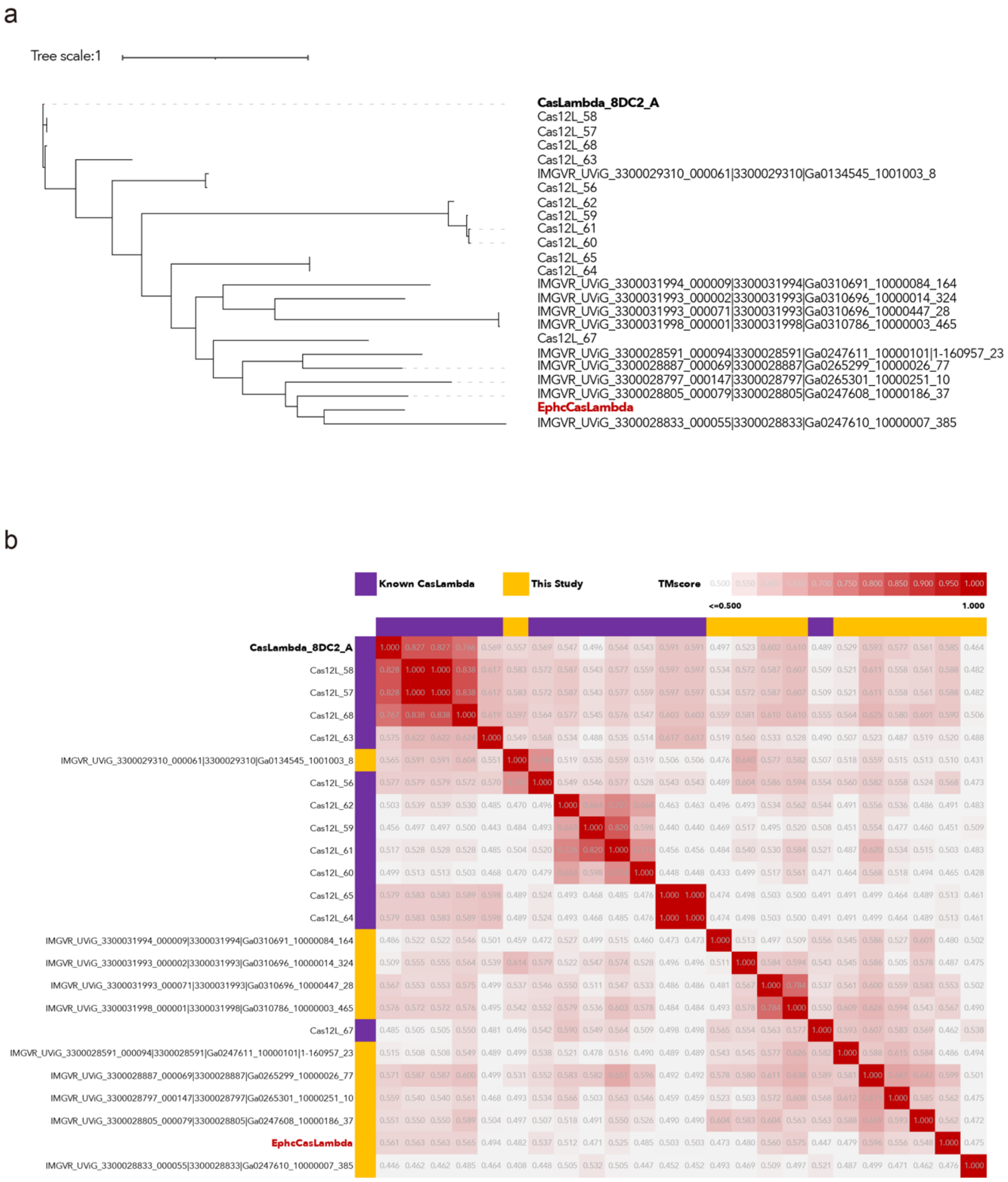


**Supplementary Fig. 5 | Phylogenetic analysis and structural alignments of the Casλ catalog. a,** Phylogenetic tree of Casλ homologs, **b**, Heatmap of TM-scores of pairwise Casλ proteins.

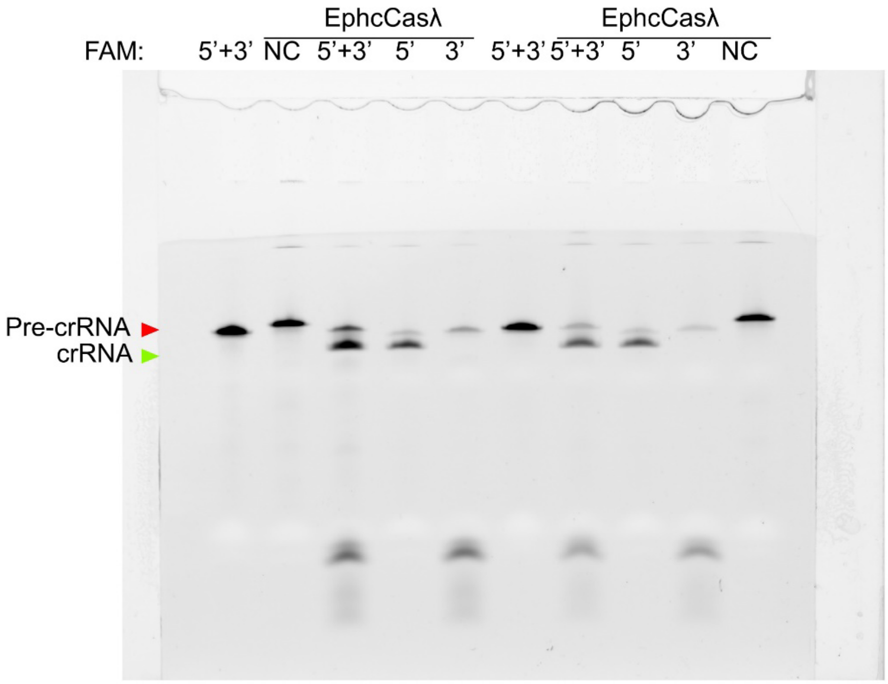


**Supplementary Fig. 6 |** Raw data for Fig. 5b. EphcCasλ specifically processes its own crRNA by cleaving several bases in the spacer region (or 3' end). Similar experiments were performed with 4 μM RNA (left lane1 to left lane5) and 2 μM RNA (left lane6 to left lane10).

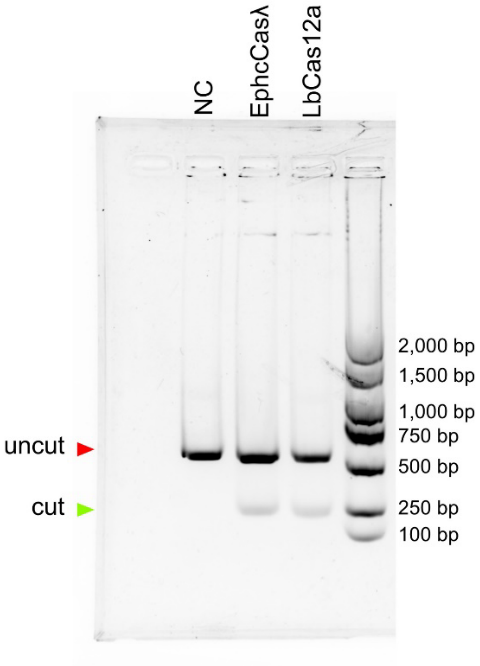


**Supplementary Fig. 7 |** Raw data for Fig. 5d.

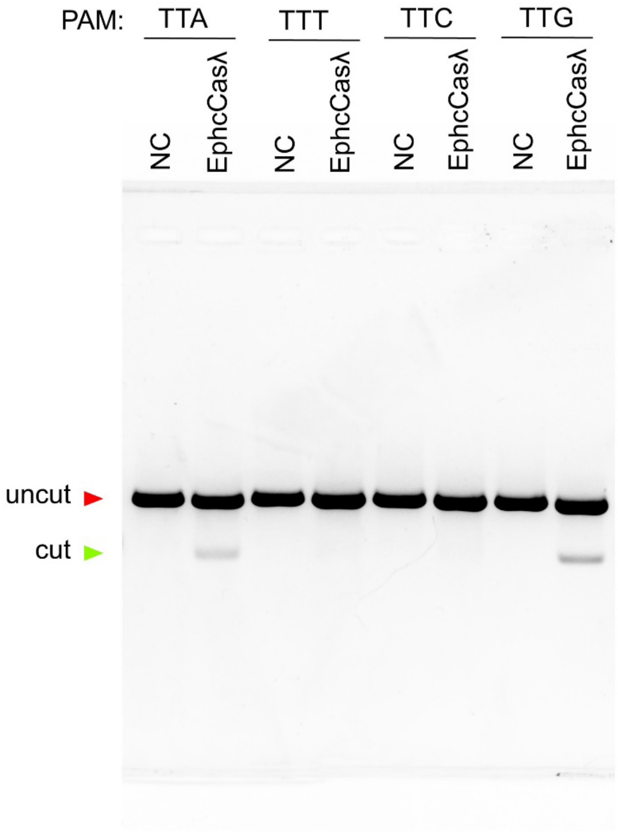


**Supplementary Fig. 8 |** Raw data for Fig. 5f.

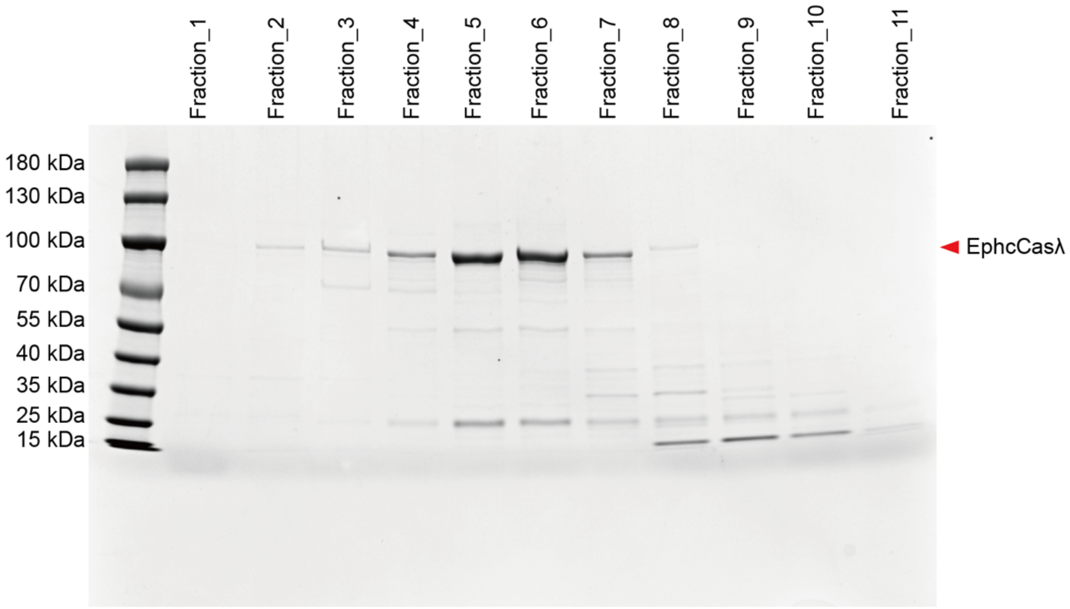

**Supplementary Fig. 9 |** Protein purification results after SEC. Sequence of EphcCasλ was optimized for *E.coli*, and has been inserted into vector pET-28a(+) with 5'-6×his and 3'-6×his, expressing in *E.coli* BL21.

**Supplementary Table 1** | Summary of nuclease-associated traits of Cas12 homologs

| **Cas12 homologs** | **Protein size** | **DNase activity** | **RNase activity** | **RNase mechanisms** | **tracrRNA-dependent** | **Self-processing pre-crRNA** | **Ref.** |
|---|---|---|---|---|---|---|---|
| Cas12a (Cpf1) | 1025 ~ 1251aa | ssDNA, dsDNA | yes | acid-base mechanism | no | **yes** | [3–6] |
| Cas12b (C2c1) | 743 ~ 1118aa | dsDNA | no | - | yes | no | [7] |
| Cas12c (C2c3) | 1209 ~ 1330aa[8] | no | yes | Metal-dependent | yes | **yes** | [9–11] |
| Cas12d (CasY) | 1080 ~ 1207aa | ssDNA, dsDNA | no | - | yes | no | [9] |
| Cas12e (CasX) | 818 ~ 986aa | dsDNA | no | - | yes | no | [12] |
| Cas12f (Cas14a) | 400 ~ 700aa[13] | ssDNA, dsDNA3/11/24 7:31:00 PM | no | - | yes | no | [13–15] |
| Cas12g | 720 ~ 830aa[8] | no | yes | Metal-dependent | yes | no | [16] |
| Cas12h | 870 ~ 924aa[8] | dsDNA | yes | Unknown | no | **yes** | [8] |
| Cas12i | 1033 ~ 1093aa[8] | dsDNA | yes | acid-base mechanism | no | **yes** | [17] |
| Cas12j (CasΦ) | 707 ~ 812aa | dsDNA | yes | Metal-dependent | no | **yes** | [18,19] |
| Cas12k (V-U5) | 393 ~ 639aa | no | no | - | yes | no | [20,21] |
| Cas12L (Casλ) | 735 ~ 828aa | dsDNA | yes | Metal-dependent | no | **yes** | [22] |
| Cas12l (Casπ) | 850 ~ 867aa | dsDNA | no | - | yes | no | [23] |
| Cas12m (V-U1) | 519 ~ 608aa | no | yes | Unknown | no | **yes** | [24,25] |
| Cas12n (V-U4) | 442 ~ 513aa | dsDNA | no | - | yes | no | [26] |
| TnpB | 372 ~ 545aa | dsDNA | yes? | Metal-dependent? | yes | no | [27,28] |

Note: The statistics of the protein sizes of Cas12 homologs were analyzed using data downloaded from CasPEDIA[29] (http://caspedia.org/).

**Supplementary Table 10** | RNA, plasmids, and primers.

| Substrate | Abrv. | Sequence | FAM | Source |
|---|---|---|---|---|
| crRNA | | | | |
| LbaCas12a crRNA target at randomized PAM dsDNA | Cas12a-cr1 | 5'-UAAUUUCUACUAAGUGUAGAUCCGGCAAGCUGCCCGUGCCC-3' | none | Synthesized by GenScript |
| EphcCasLambda crRNA target at randomized PAM dsDNA | crRNA | 5'-AUUGUUGGAAUAUCACUUUUGUAGGGUAUUCACAACCCGGCAAGCUGCCCGUGCCC-3' | none | Synthesized by GenScript |
| EphcCasLambda crRNA with both end FAM target at randomized PAM dsDNA | crRNA_5/3 | 5'-AUUGUUGGAAUAUCACUUUUGUAGGGUAUUCACAACCCGGCAAGCUGCCCGUGCCC-3' | 5'+3' | Synthesized by GenScript |
| EphcCasLambda crRNA with 5' end FAM target at | crRNA_5 | 5'-AUUGUUGGAAUAUCACUUUUGUAGGGUAUUCACAACCCGGCAAGCUGCCCGUGCCC-3' | 5' | Synthesized by GenScript |

| | | | | |
|---|---|---|---|---|
| randomized PAM dsDNA | | | | |
| EphcCasLambda crRNA with 3' end FAM target at randomized PAM dsDNA | crRNA_3 | 5'-<br>AUUGUUGGAAUAUCACUUUUGUAGG<br>GUAUUCACAACCCGGCAAGCUGCCC<br>GUGCCC-3' | 3' | Synthesized by GenScript |
| Another crRNA with both end FAM target at randomized PAM dsDNA | ncrRNA | 5'-<br>CUCGCGGUCCCAUCGGAACGGGUUG<br>UGGUUCCGACCCGGCAAGCUGCCCG<br>UGCCC-3' | 5'+3' | Synthesized by GenScript |
| Plasmids | | | | |
| optimized DNA sequence of EphcCasLambda for *E. coli* inserted into vector pET-28a(+) | pET-28a(+)_EphcCasLambda | 5'-<br>atggctcaacacaaatcaaataacgaggaaagtgcgattaat<br>aagactttcattttcaaagcgaagtgcgataaaaacgatgtaat<br>cagcctgtgggaaccggcggcaaaagaatactgcgattact<br>ataacaaggtgagcaaatggattgctgacaacctgatcaccat<br>gaagatcggtgacctggctcagtatatcacgaatcagaactc<br>caagtattataccgcggttacaaacaagaaaaagaaggatctt<br>ccgctgtaccgcatttttcagaagggtttttcctcgcaatgtgca<br>gacaacgcgttgtactgcgcgattaagagcatcaacccggaa<br>aattataaaggcaactcgctgggcattggcgaaagcgactac<br>cgtcgtttcggctacatccaatccgtggtgtccaacttccgcac<br>caaaatgagcagcttgaaggcgaccgttaaatggaaaaagtt<br>cgatgtgaacaatgtggacgacgagaccttgaaaatccaaac<br>catctatgacgtcgacaaatatggtatcgaaactgccaaggaa<br>tttaaagagctgatcgaaaccttgaagacccgtgttgaaacgc<br>cacagctgaacgataccattgcccgtctggagtgcctttgcga<br>ctactacagcaagaacgagaaggcgattaacaacgaaattg<br>aaacaatggcaattgcggatctgcagaagtttggtggctgtca<br>gcgtaagtccttgaatgccttcactatccacaagcaggatagc<br>ctcatggaaaaggtgggcaacaccagctttcgcctgcaactg<br>ccgtttagaaaaaagacgtacgtgatcaatctgctgggtaacc<br>gtcaggttgtgaacttcgtgaacggtaaacgtgttgatttgattg<br>acatcgcggaaaatcatggtgacttggttacgttcaacatcaa<br>aaatggtgttctcttcgtgcacctgaccagcccgattgtgttcg<br>ataaagacgttcgggatattcgtaatgtggtgggtatcgacgt<br>gaacatcaagcactccatgctggcaacctccatcaaagatgt<br>gggtaatgttaaaggctatatcaacttgtacaaagagttgttga<br>acgatgatgagttcgttagcacctgtaatgagtccgaactggc<br>gctgtaccgtcaaatgagcgagaatgttaatttcggcatccta<br>gagactgactctttgtttgaacgcatcgtgaatcagagcaaag<br>gtggctgcttaaagaataaactgatccgccgcgagctcgctat<br>gcagaaagtcttcgagcgcatcaccaaaacgaataaggatc<br>agaacattgttgattatgtgaactacgtcaaaatgatgcgtgcg<br>aaatgcaaagcgagctatattctgaaagagaaatacgatgag<br>aagcaaaaagaatactatgtgaagatgggttttaccgacgagt<br>ctaccgaaagcaaggaaacgatggacaaacgtcgcgaaga<br>gttcccgtttgttaataccgatacggccaaagagctgctggtg<br>aagcaaaacaacattcgtcaagatatcattggctgtcgtgaca<br>acatcgtcacctatgcattcaatgtcttcaagaataacgaatac<br>gacaccctgtccgttgaatacctggatagtagccaatttgataa<br>gcgtcgtattgcaaccccgaaatctctgcttaagtaccataaat<br>tcgagggcaagactaaggacgaagttgagaacatgatgaaa<br>agcgaaaaactgtcgaatgcgtattacaccttcaagtacgaaa<br>acgacgttgtttccgacattgattacagcgacgagggtaacct<br>gcgccgtagcaagttgaattttggtaattggattattaaatctat | | Synthesized by GenScript |

| | | | | |
|---|---|---|---|---|
| | | ccactttgcagacattaaggacaaatttgttcagctgtctaacaacaataaaatgaatatcgtgttctgcccgagcgcgtttagctcgcaaatggattctattacccacaccctgtattacgttgaaaagatcaccaaaaacaaaaagggcaaagagaagaagaaatacgtactagctaataagaaaatggtccgtacgcagcaggagaaacatatcaacggcctgaacgctgattataactcagcgtgcaatctgaagtacatcgctctgaacgacgagctgcgcgacaaaatgaccgatcgttttaaggcgagcaaaaagattaagaccatgtataacatcccggcgtataatatcaagagcaacttcaagaaaaacctgtctgcgaaaaccatccaaacctttcgcgagttaggtcattatagagacggtaaaattaacgaggacggcatgtttgtagagaacttggag-3’ | | |
| Synthetic target dsDNA sequence with 5’-TTA PAM insert into vector pUC-19 | pUC-19_TTA | 5’-cagccaagcttgcatgcctcgtgacctgcgctttagtttttttaccggcaagctgcccgtgcccctcccgaattcccggtctccctatagtgagtcaggcgg-3’ | | Synthesized by GenScript |
| Synthetic target dsDNA sequence with 5’-TTT PAM insert into vector pUC-19 | pUC-19_TTT | 5’-cagccaagcttgcatgcctcgtgacctgcgctttagttttttttccggcaagctgcccgtgcccctcccgaattcccggtctccctatagtgagtcaggcgg-3’ | | Synthesized by GenScript |
| Synthetic target dsDNA sequence with 5’-TTC PAM insert into vector pUC-19 | pUC-19_TTC | 5’-cagccaagcttgcatgcctcgtgacctgcgctttagtttttttcccggcaagctgcccgtgcccctcccgaattcccggtctccctatagtgagtcaggcgg-3’ | | Synthesized by GenScript |
| Synthetic target dsDNA sequence with 5’-TTG PAM insert into vector pUC-19 | pUC-19_TTG | 5’-cagccaagcttgcatgcctcgtgacctgcgctttagtttttttgccggcaagctgcccgtgcccctcccgaattcccggtctccctatagtgagtcaggcgg-3’ | | Synthesized by GenScript |
| Plasmid with randomized PAM sequence | p11-3’_8xN_PAM-site1 | | | addgene cat.no.160132 |
| Primers | | | | |
| Primers for preparing randomized PAM dsDNA substrate | PAM_F1 | 5’-GTTCTTTCCTGCGTTATCCCCT-3’ | | Synthesized by GenScript |
| | PAM_R1 | 5’-CCCAGTCTTTCGACTGAGCCTT-3’ | | Synthesized by GenScript |
| Primers for capturing and enriching uncleaved dsDNA | PAM_F2 | 5’-CGCCGCAGCCGAACGACC-3’ | | Synthesized by GenScript |
| | PAM_R2 | 5’-CCGCCAAAACAGCCAAGC-3’ | | Synthesized by GenScript |
| Primers for preparing TTA/TTT/TTC/TTG dsDNA substrate | TTN_F1 | 5’-GCGTTTCGGTGATGAC-3’ | | Synthesized by GenScript |
| | TTN_R1 | 5’- CGGCCTTTTTACGGTTCCT-3’ | | Synthesized by GenScript |
